# Insect Wing Classification of Mosquitoes and Bees Using CO1 Image Recognition

**DOI:** 10.1101/034819

**Authors:** Nayna Vyas-Patel, Sai Ravela, Agenor Mafra-Neto, John D Mumford

## Abstract

The certainty that a species is accurately identified is the cornerstone of appearance based classification; however the methods used in classical taxonomy have yet to fully catch up with the digital age. Recognising this, the CO1 algorithm presented on the StripeSpotter platform was used to identify different species and sexes of mosquito wings (Diptera: Culicidae) and honey bee and bumblebee wings (Hymenoptera: Apidae). Images of different species of mosquito and bee wings were uploaded onto the CO1 database and test wing images were analysed to determine if this resulted in the correct species being identified. Out of a database containing 925 mosquito and bee wing images, the CO1 algorithm correctly identified species and sexes of test wing image presented, with a high degree of accuracy (80% to 100% depending on the species and database used, excluding sibling species) highlighting the usefulness of CO1 in identifying medically important as well as beneficial insect species. Using a larger database of wing images resulted in significantly higher numbers of test images being correctly identified than using a smaller database. The hind wings of Hymenoptera provided higher levels of correctly identified results than using the fore wings. The software should be used in conjunction with other identifying criteria (salient morphological features) in addition to the wings. CO1 is a powerful algorithm to use in identifying insect wings in its current form and has great potential if it is adapted and tailored for insect species identification. It is suggested that a primary aim in the digital age should be the production of a ‘World Wide Database’ of insect images, where all known insect images can be made available to everyone, with image recognition and species knowledge at its core.

## Introduction

The distinctive pattern of veins on winged insects is characteristic of the species and affords the opportunity to separate insect species using image recognition software. Previous attempts at identifying insect species using automated, semi-automated and geometric morphometric methods (ABIS, Draw wing, tpsdig2, MOBS and API class) to distinguish between insect wing images, have been comprehensively described by Hall (2011). Few insect identification web sites utilise modern image recognition software to aid the identification process. CO1 image recognition is one such method and its potential to identify insect species using wing images was explored.

A body of work exists describing a number of different technological methods used to identify insect species by their wing venation. Jing Dai (2006) and Zhou et al (1985) utilised computer aided pattern recognition and digitized wing images to distinguish between insect species. Closer to the image recognition idea, Lamprecht (2010) worked on Shutterbug, photographing entire insects rather than just the wings from many different angles and under differently coloured lights, for use in image recognition software. Using the entire insect body, Yang et al (2010) extracted 14 features such as insect sphericity, elongation, also rectangularity and used an algorithm called Random Trees to identify insects. An Lu (2010), again using the entire insect body, created rows and columns of images but limited in number (Sparse Representation) of the whole body, head, thorax, abdomen and wings to distinguish between species. Zhao et al (2009) reported an image recognition method which used clustering of insect pests of sugarcane. Wang et al (2012) used Artificial Neural Networks and a Support Vector Machine as pattern recognition methods to identify insects to the order level. Al-Saqer et al (2011) described an image processing system for identifying Pecan weevils. Zhang et al (2013) described high resolution electronic image pre-processing for the identification of stored grain insects. This plethora of interest in image recognition of insects attests to the importance and the varied applications of this field of research. A number of insects are pests of food (weevils) or vectors of disease (mosquitoes), whilst other species such as bees are invaluable in pollination and thus the production of food. Put simply, it is vital that easy to use, free and accurate methods are created to identify insect species.

The methods above varied in the technology and approach used. They are different to the more recent image recognition methods available currently which are freely available online and easy to use. Typical image recognition systems use optical character recognition and are computerised methods of examining an image; comparing it with other images in the database and identifying it. This study examined the use of CO1, a freely accessed image recognition algorithm (Lahiri et al 2011a & b), presented as one of the options on the StripeSpotter Platform. The other option on StripeSpotter - the Stripe code was developed for recognising stripes and spots on zebras and leopards. CO1, not being not limited to the recognition of any one particular shape or form, was used here.

The CO1 algorithm was developed to identify individual salamanders, more specifically, the distinctive patterns on the backs of a threatened species of marbled salamander – *Ambystoma opacum* (Ravela & Gamble, 2004). It creates overlapping multi-scale differential features along the entire length of an image; these features are then composed into histograms (Ravela & Gamble, 2004 & Lahiri et al 2011a). The string of multi-scale histograms is treated as a vector and correlated to deduce similarity. A database of images is created by the user containing a number of stock images of the organism/s in question. The query image and its histogram vector can then be automatically compared with each database histogram vector and the corresponding images (the results) are ranked by their score within a matter of seconds. Each ranked image is automatically given a score which appears as the ‘cost’ value of each of the ranked results. The closer the cost value is to 1, the more certain it is that the test image is a good match of the rank 1 image that is retrieved by the software from the database. This ‘cost’ value falls incrementally for every image further down the ranks, only the rank 1 cost values were considered here. The rank 1 cost values of images which had/did not have an identical copy in the database were noted. Wing images which had no sample image of the test species in the database were also considered. Cost values of totally alien, non-wing images, which were not in the database, were examined. If a large number of images were present in the database, a maximum of 100 possible matches were ranked, with the closest matches being rank 1 or near to rank 1 (Lahiri et al 2011a & b). The certainty that the correct species was being identified could be substantiated if the images retrieved further down the ranks were examined and found to be the correct species. Hence the totals of accurately identified images up to rank 5 and rank 10 were also noted, in addition to rank 1 and rank 2 images.

The primary aim of the study was to distinguish between different species using CO1 (the freely available version on the StripeSpotter platform). The ability of the algorithm to distinguish between sibling species was examined in one case (*Bombus* sub species), but this was not the main aim. The intention was to test the algorithm in ‘real use’ scenarios, where equal numbers of each species, the use of sophisticated cameras/equipment was not possible. This would indicate the accuracy of the software in the hands of the citizen scientist, where numbers of available specimens, sophisticated equipment, or any kind of specialized laboratory condition was realistically not the norm. StripeSpotter automatically describes how good/bad every image is (good, bad, or ok) and together with the human eye it is possible to create good quality images easily. A further study using 12 good quality images (6 male, 6 female) of each species in the database was used to ascertain if this would yield more accurate results. The results from the larger and smaller databases were examined to determine if database scale had an effect on accuracy. CO1 was also tested for its ability to separate the sexes. A third database with 30 images of each species was also tested, this time with test wing images which did not have the exact copy in the database. The optimum use of the software in order to arrive at accurate species identification was discussed.

## Materials and Methods

Insect specimens were obtained from colonies of bumblebees reared by Royal Holloway College, UK (GPS: 51.426347, −0.562731), honey bees reared by Surrey Beekeepers UK (GPS: 51.401348, −0.259752) and laboratory reared mosquitoes from the London School of Hygiene and Tropical Medicine UK (GPS: 51.520707, −0.129994). The wings were dissected out from the body under a standard dissection microscope and photographed with a Samsung NV10 digital camera, using only the sub-stage lighting of the microscope as this produced a clear image of the wing shape and venation. Each image was uploaded into an Adobe Photoshop (CS5) image editor and rotated so that the point of insertion of the wing into the body of the insect (known as the Jugum) always faced to the left and the wing was aligned to be as horizontal as possible, using Image Rotate in the top menu bar of Photoshop as depicted in Figure 1 & 2. The newly aligned and rotated images were saved as .jpg files, creating a different file for each species, sex and where appropriate, fore and hind wings.

**Figure 1.**
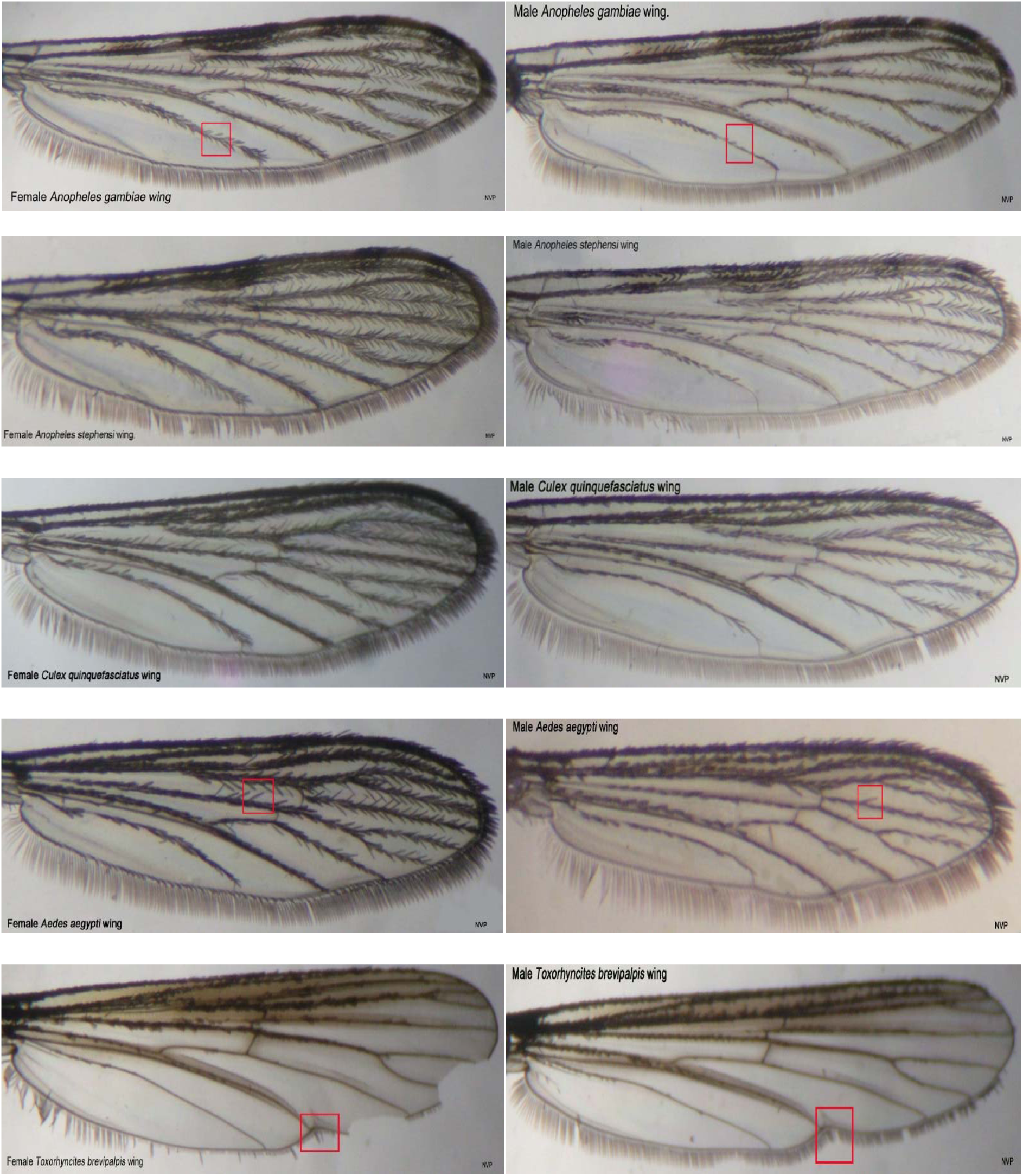
Images of Female and Male Mosquitoes (Diptera: Culicidae).

**Figure 2.**
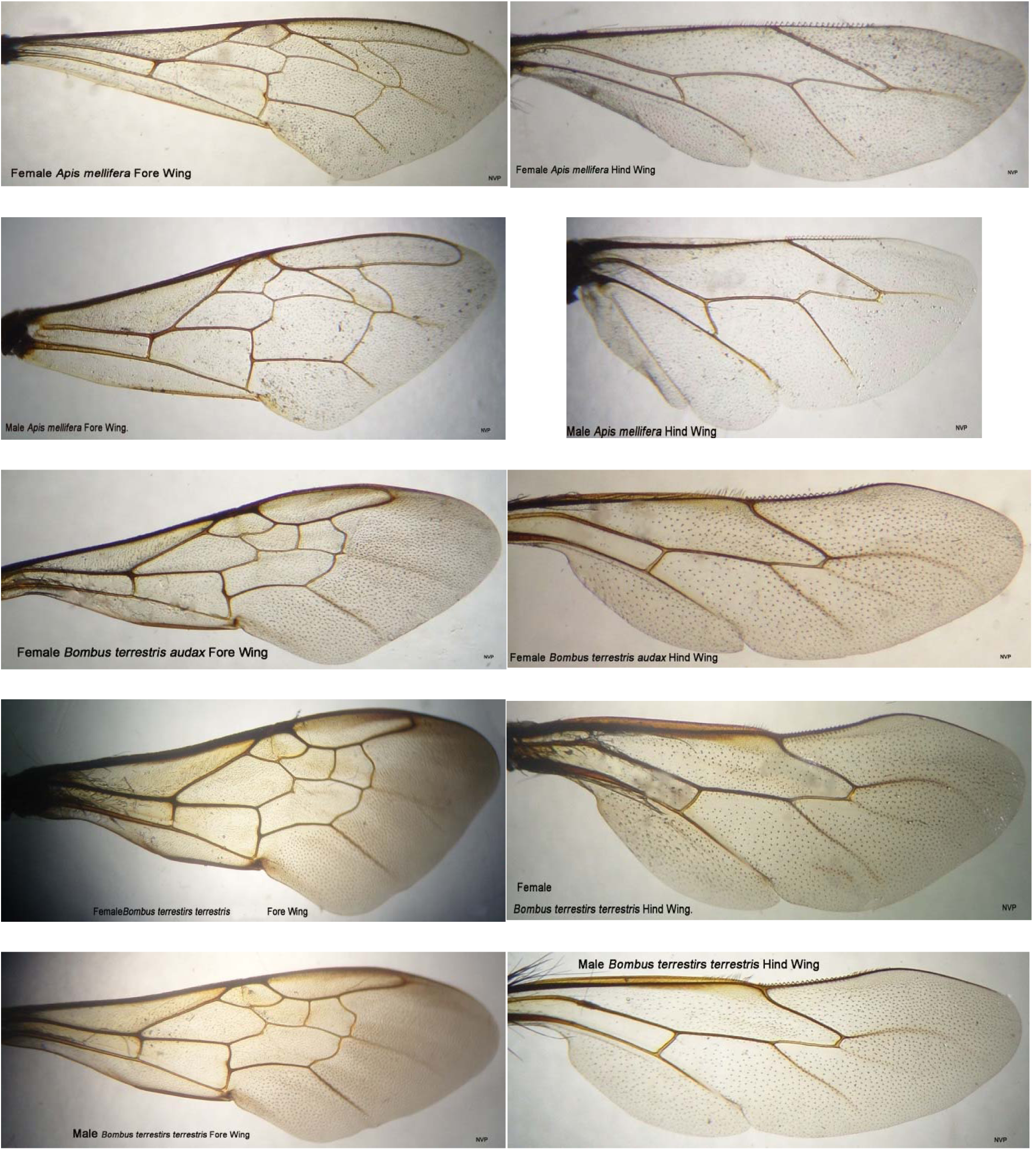
Images of the Fore and Hind Wings of Female and Male Hymenoptera.

StripeSpotter was downloaded from the internet and images of insect wings were uploaded into the database. Of the mosquitoes (Diptera: Culicidae), a total of 59 *Anopheles gambiae*, 113 *Anopheles stephensi*, 244 *Culex quinquefasciatus*, 140 *Aedes aegypti* and 29 *Toxorhyncites brevipalpis* wings were uploaded into the database. Of the honey bees and bumblebees (Hymenoptera: Apidae), a total of 40 fore wings and 57 hind wings of female *Apis mellifera;* 80 fore wings and 30 hind wings of male *Apis mellifera* (honeybees); 33 fore and 24 hind wings of female bumblebee workers *Bombus terrestris audax* (Bta); 43 fore and 33 hind wings of male *Bombus terrestris terrestris* (Btt) were uploaded. In all, a grand total of 925 wing images of both Dipteran and Hymenopteran wings were uploaded into the Large Database (LDB). 50 images each of *Anopheles gambiae, Anopheles stephensi, Aedes aegypti, Culex quinquefasciatus, Apis mellifera* and 30 images of *Toxorhyncites brevipalpis* and *Bombus* sub species, were analysed using the large database set, containing a total of 925 images of both Dipteran and Hymenopteran wings and using the CO1 algorithm (Table 1).

**Table 1.**
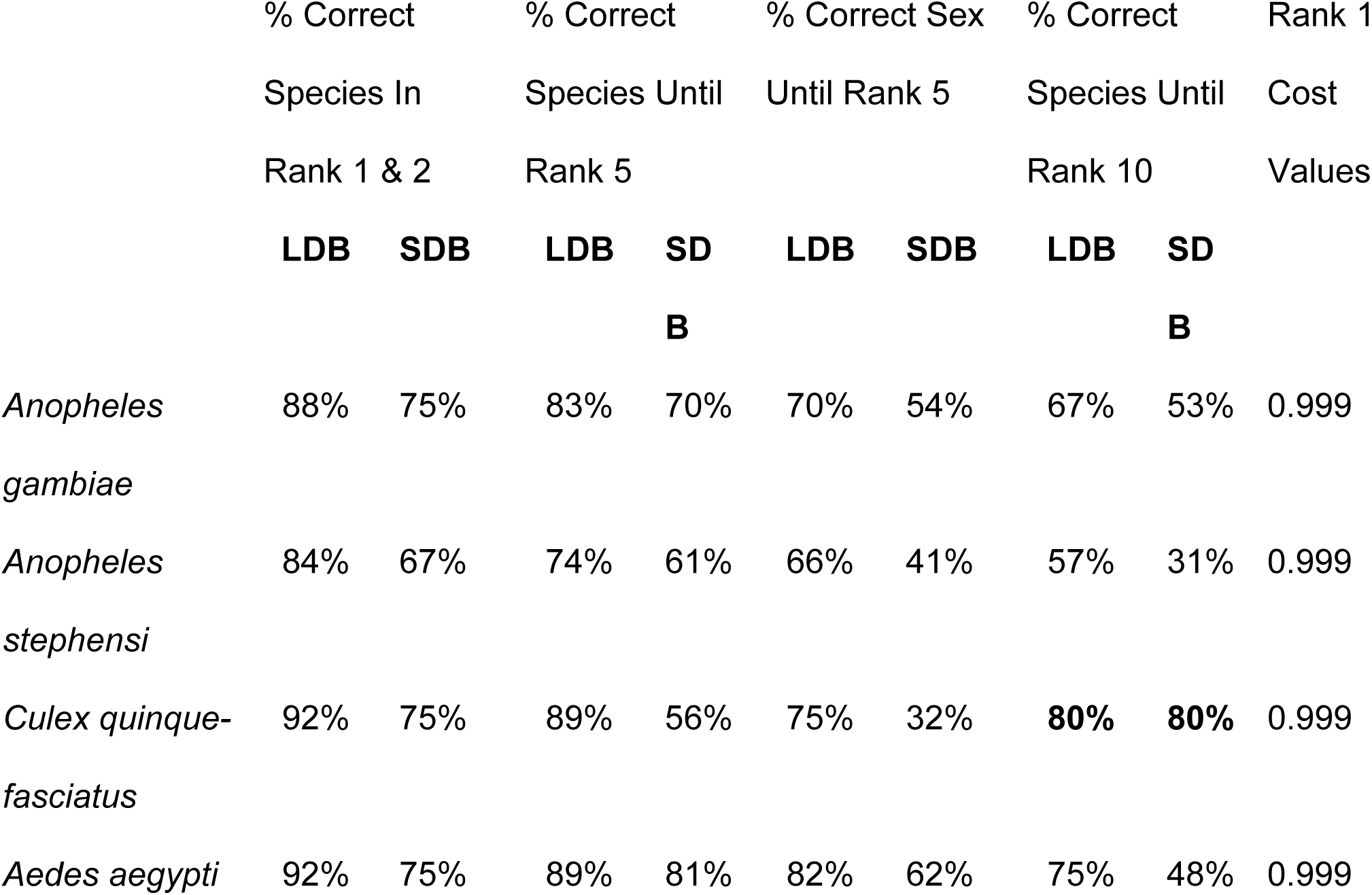
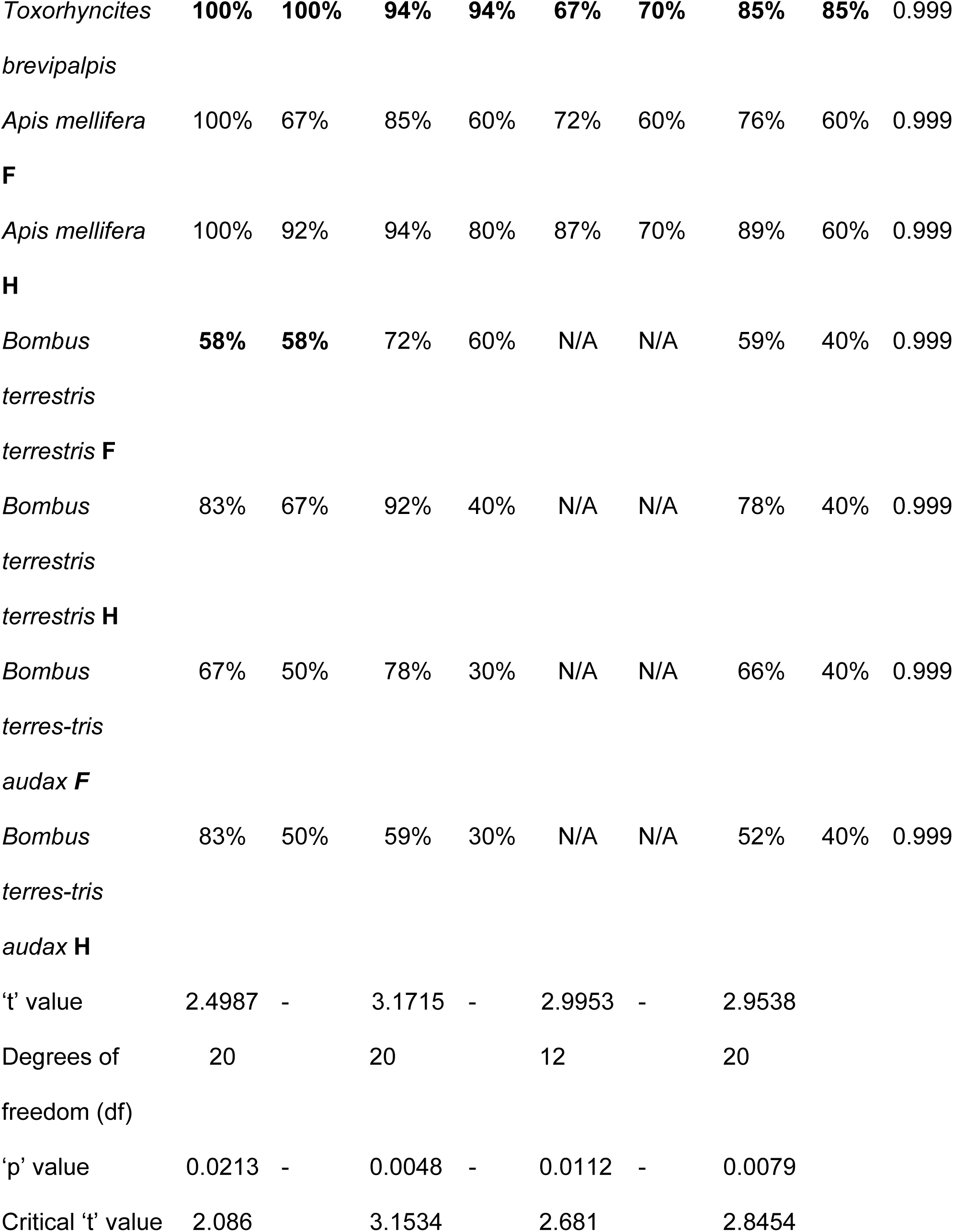
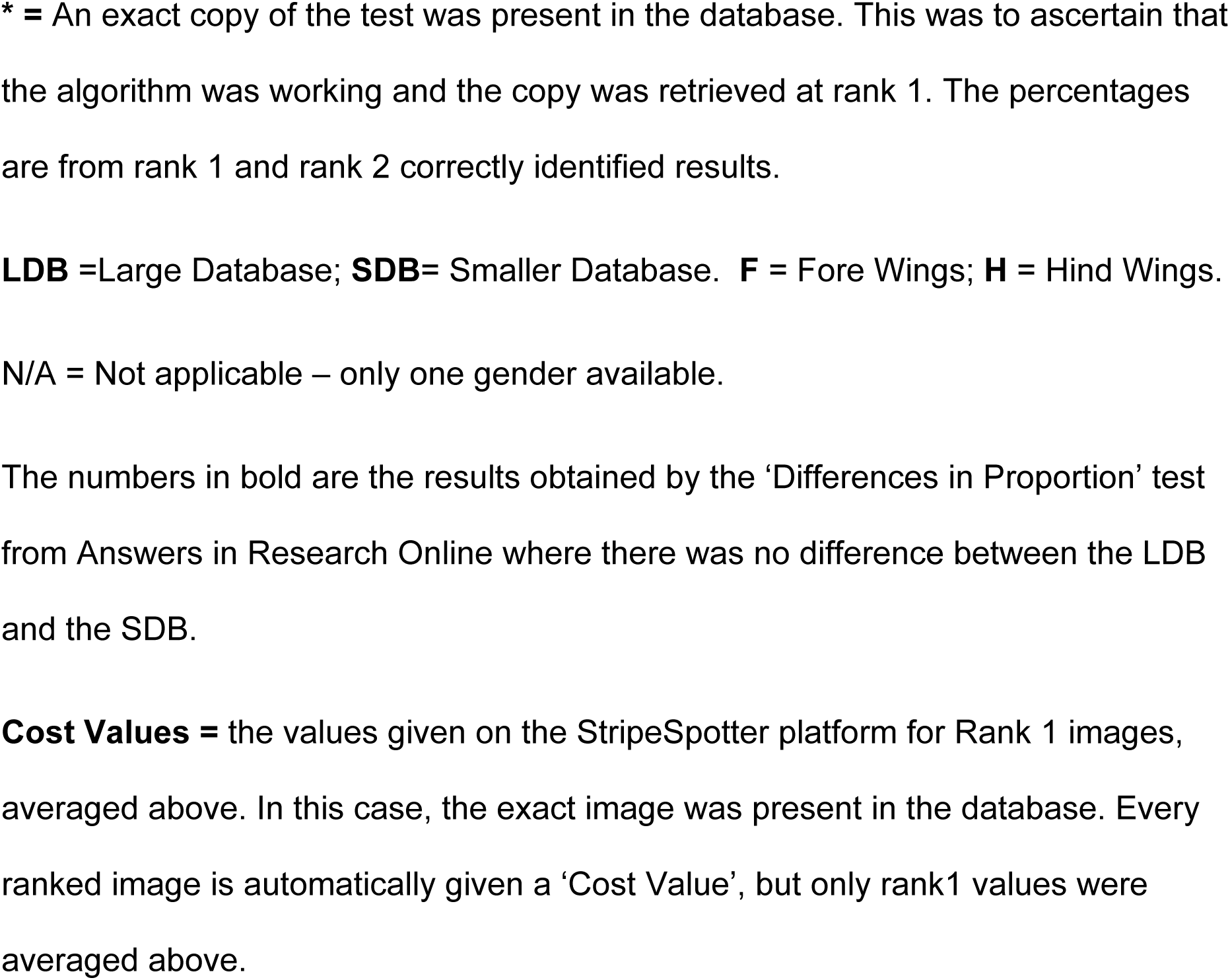
Percentage of Correctly Identified Diptera and Hymenoptera Wings*.

To determine if the software worked to retrieve the exact image when tested with images that were in the database, images already in the database were used as the test and the results of rank 1 and 2 noted. This is a vital first step when using any new software as it may not retrieve exact copies of a test image if present in a database, indicating that software improvements were required. To ensure that this did not affect the results, a second test was performed with new images that had no copies in the database, but which were from the same species as those in the database. The results were compared for both sets (Table 1a).

**Table 1a:**
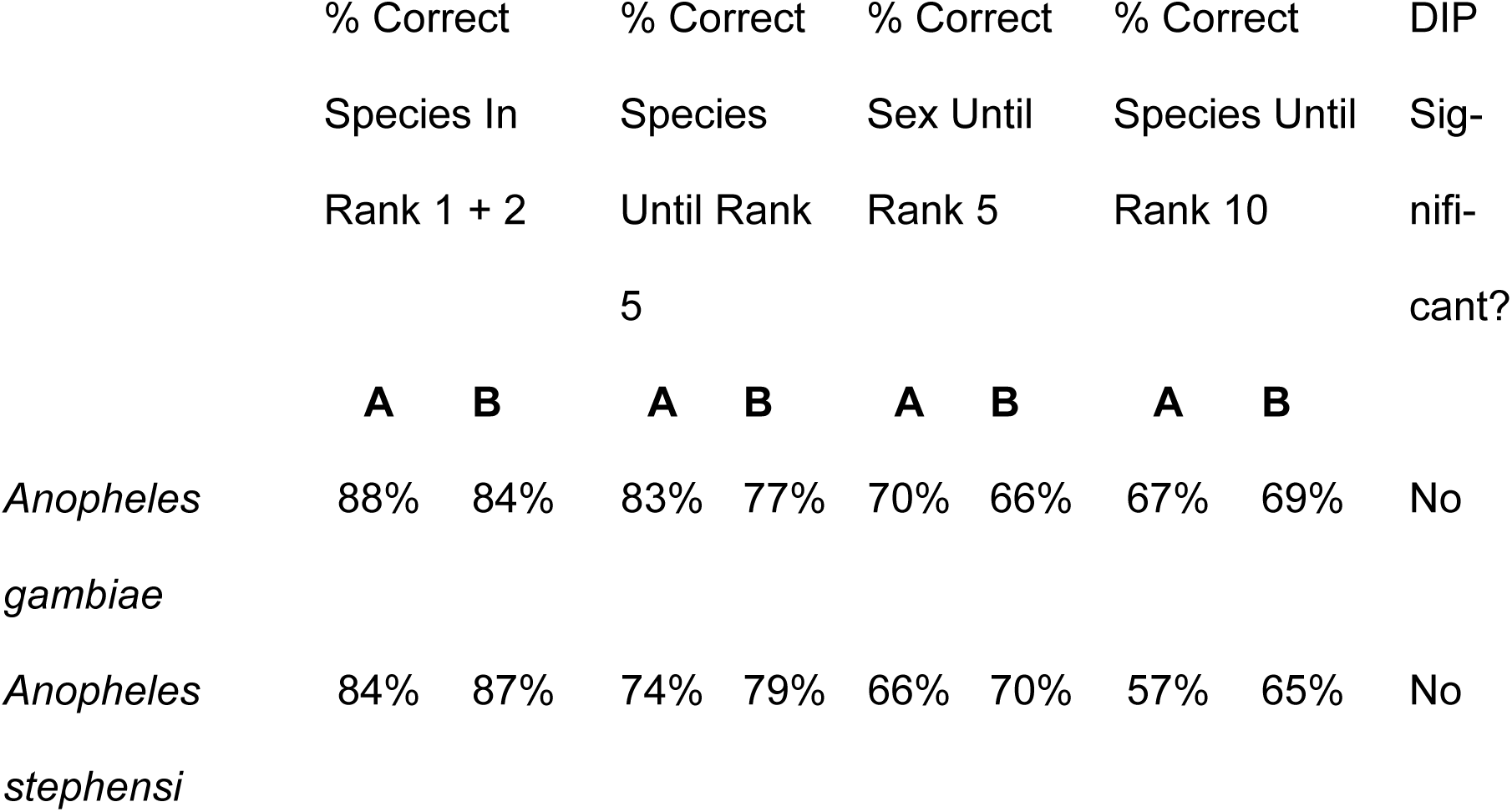
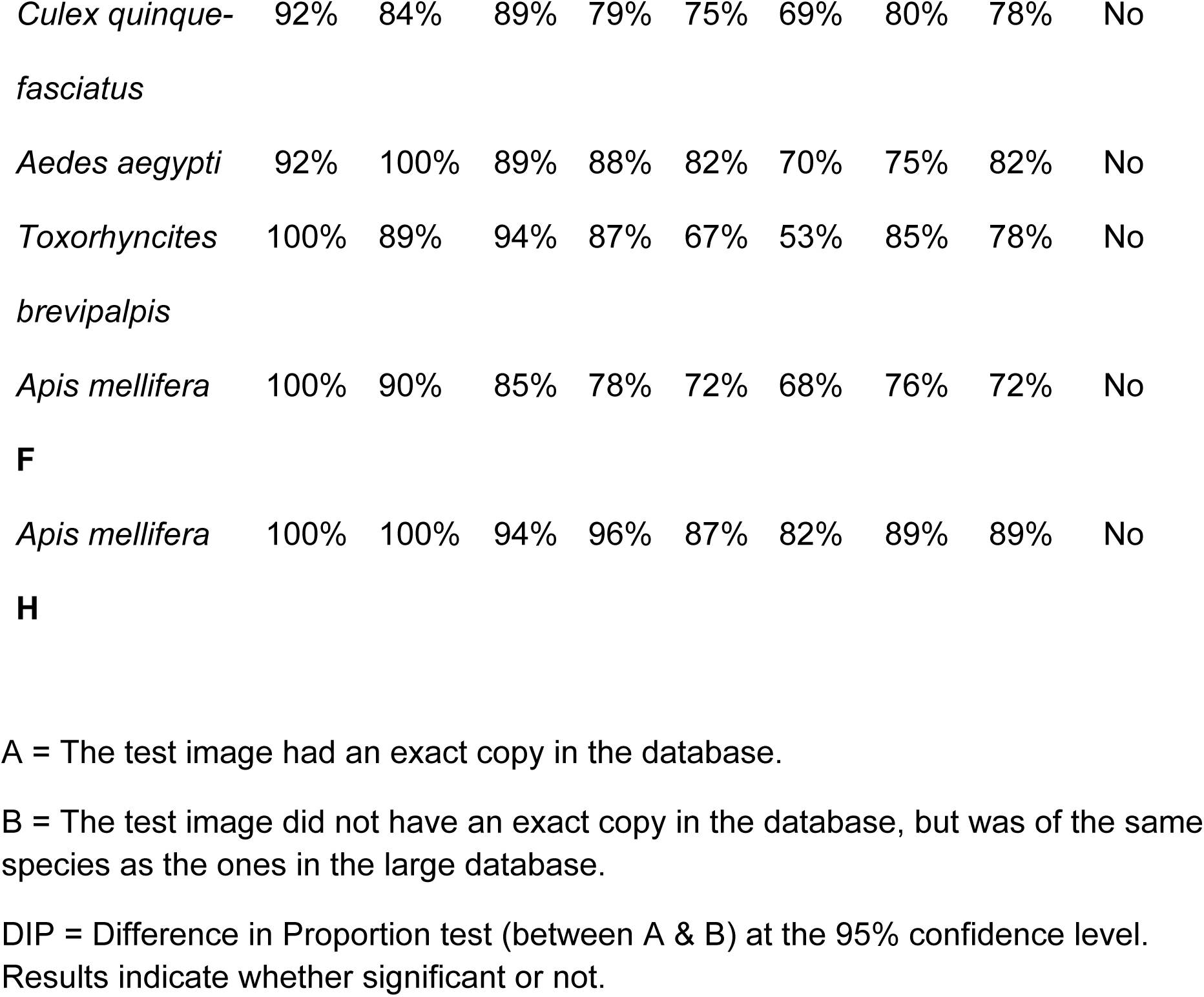
Comparison of results Using Test images, Exact Copies of which Are (A), or Are Not (B), in the Large Database.

The Smaller Database (SDB) contained 12 images from each species (6 images of each sex where applicable). Furthermore, in the case of the bees and bumblebees (Hymenoptera), 6 images each of the fore and hind wings of each species were uploaded. A total of 108 images were uploaded into the Smaller Database (SDB). The median scores of the larger and smaller databases were compared, Table2. A third database was created which had 30 good quality (‘good’ as defined by StripeSpotter where every image is assessed to be good, bad or ok) images, 15 male, 15 female, of each of the species. This was tested with 50 new wing images (25 in the case of *T. brevipalpis* and *Bombus* sub species), a copy of which was NOT in the database, Table 3.

**Table 2.** The Values for the Median and the Mode (Diptera).

|  | Median (from<br>totals up to<br>Rank 5) |  | Mode (from<br>totals up to<br>Rank 5) |  | Median (from<br>totals up to<br>Rank 10) |  | Mode (from<br>totals up to<br>Rank 10) |  |
| --- | --- | --- | --- | --- | --- | --- | --- | --- |
|  | **LDB** | **SDB** | **LDB** | **SDB** | **LDB** | **SDB** | **LDB** | **SDB** |
| *Anopheles*<br>*gambiae* | 4 | 4 | 5 | 4 | 7 | 5 | 8 | 5 |
| *Anopheles*<br>*stephensi* | 4 | 3 | 4 | 3 | 6 | 6 | 6 | 6 |
| *Culex* | 5 | 3 | 5 | 4 | 8 | 5 | 8 | 5 |
| *quinquefasciatus* |  |  |  |  |  |  |  |  |
| *Aedes aegypti* | 5 | 3 | 5 | 3 | 8 | 4 | 8 | 4 |
| *Toxorhyncites* | 5 | 5 | 5 | 5 | 9 | 7 | 9 | 8 |
| *brevipalpis* |  |  |  |  |  |  |  |  |

Values calculated from the total numbers of correctly identified species up to Rank 5 and Rank 10. **LDB** = Large Data Base. **SDB** = Small Data Base.

**Table 3.**
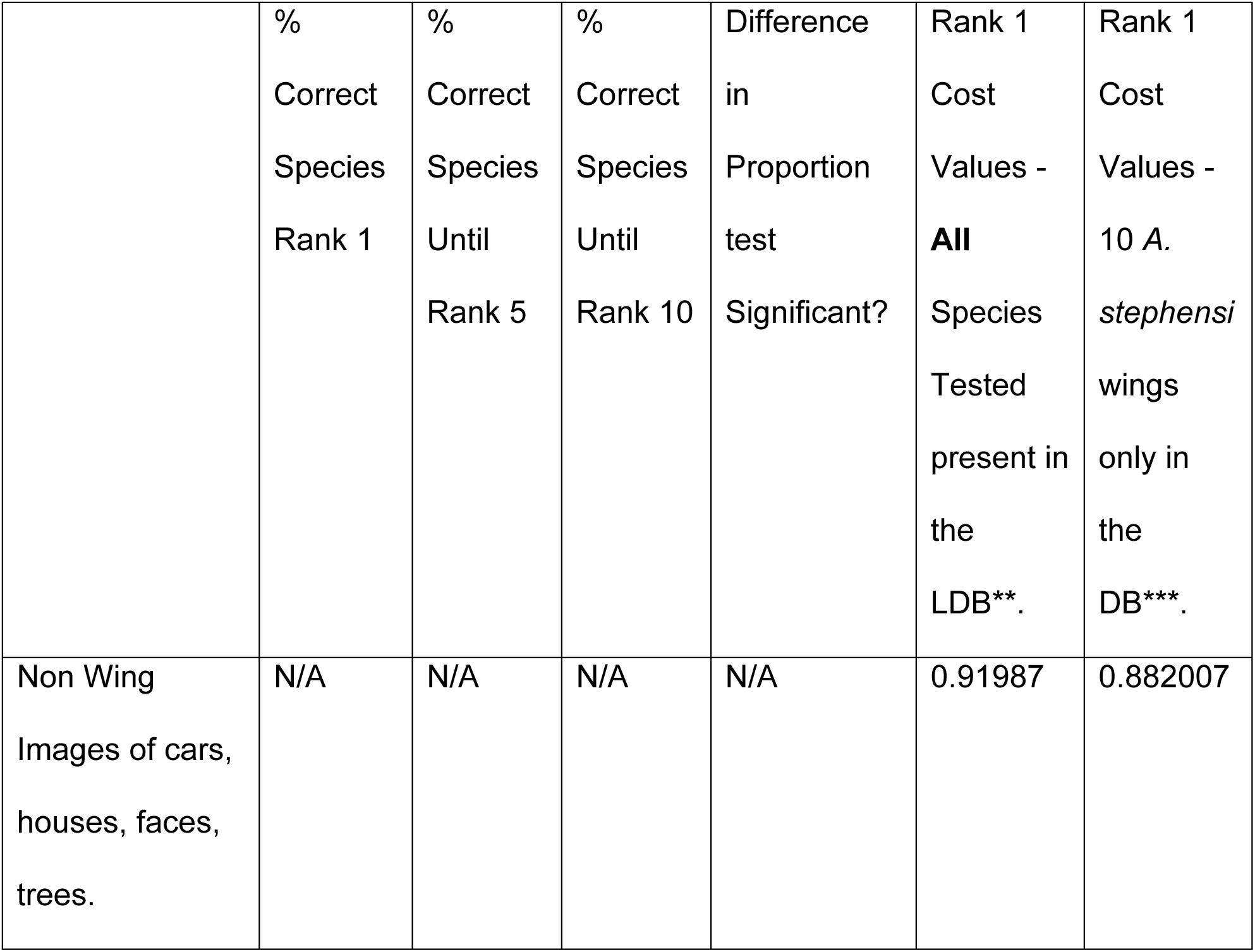
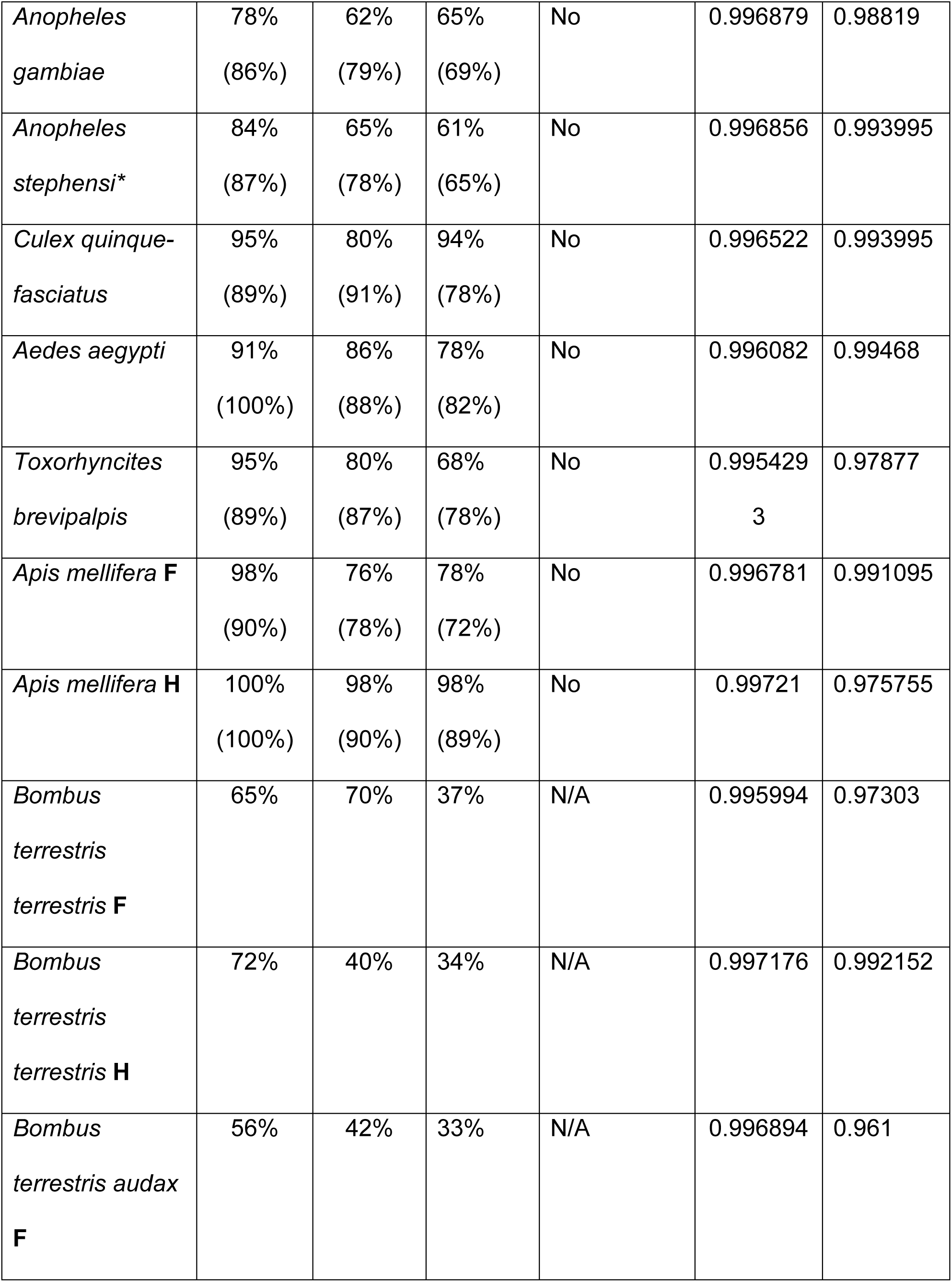
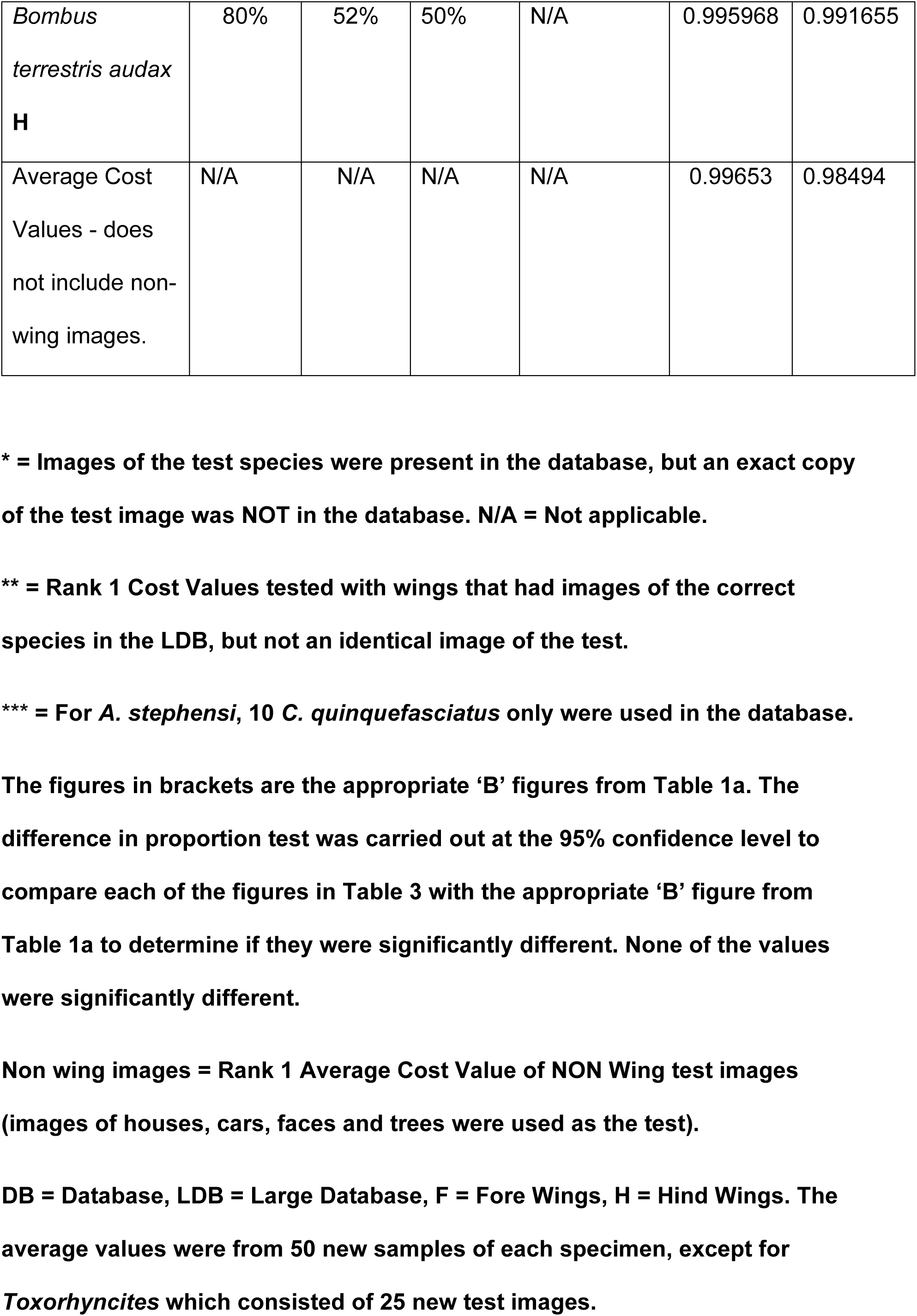
Percentage of Correctly Identified Wings^*^ and Average Cost Values, using the 3^rd^ Database.

Wing images were then tested on the StripeSpotter platform using CO1, to determine if the correct species and sex was identified, firstly within the first two, then first five ranks of results and subsequently in the first ten ranks. The results were noted for each rank, the online Graph Pad student’s ‘t’ test was used to calculate the scores as well as a ‘Difference in Proportion’ test (from ‘Answers in Research’ online).

The ‘Cost’ values of rank 1 results were noted (Table 4) for each of the following conditions:

1) When an exact copy of the test wing image was present in the database (it should be retrieved at rank1 if the algorithm performed accurately). The large database, LDB, was used and cost values of the retrieved images at rank 1 were noted.
2) When an exact copy of the test wing image was not present in the database, but other images of the test species were. The third database was used and cost values noted for rank 1 images retrieved.
3) When no images of the test wing species were present in the database. A database comprising of 10 *A. stephensi* wings only (5 of each sex) was used. For testing *A. stephensi* wings, a database of 10 images of *C. quinquefasciatus* wings (5 of each sex) was used.
4) When NON wing images were used as the test image. Images of cars, houses, trees and faces were tested using the large database (LDB) and the database containing only 1 species of wings (*A. stephensi* wings only). These images would be totally different from any other image in the database, as the database only contained images of wings.

**Table 4:** A Summary of the Cost Values of Images retrieved at Rank 1 from Different Databases & Different Test Images.

| The different Testing Regimes | Average Cost Value at Rank 1 |
| --- | --- |
| A copy of the test image was present in the LDB. | 0.999 |
| No Copies of the test image in the LDB, however other wing images of the test species were in the LDB. | 0.99653 |
| The test had no images of its own species in the DB. | 0.98494 |
| Totally alien, Non-Wing images used as the test image. | 0.91987 |

DB= Database; LDB = Large Database.

The average cost values under the different testing conditions were summarised (Table 4).

## Results

Analysis of Mosquito (Diptera: Culicidae) Wings using the Larger and Smaller Databases.

## Discussion

High levels of accurate identifications at rank 1 and 2 (Table 1, 1a & 3) were achieved. The results ranged from 80% to 100% for the large database (LDB) and from 62% to 92% for the smaller database (SDB), not taking into account the broken wings of *T. brevipalpis* where there were 95% and 100% of wings accurately identified at rank 1, but the cost values were lower and the subspecies of *Bombus*, Btt (native to Europe) and Bta (native to the U.K.) (Table 1 & 3), where the values were also lower. *A. mellifera* fore and hind wings were recognised 100% of the time early on in the ranks 1&2, unlike the sub species of *Bombus* whose fore wing scores were lower. This was because the subspecies Btt was identified as Bta and vice versa and was not due to any misidentification with other species. The high degree of accuracy of *T. brevipalpis* recognition may be due to the fact that this is one of the largest species of mosquito known, possessing much larger wings; some of the wings were broken at the edges making them distinctive; furthermore the wings have a kink, not present in the other species (Figure 1, red box).

To ensure that these results were not a consequence of using as test an image which was already in the database, new images (exact copies of which were NOT in the database) were used as test images for the large database (LDB) and the results compared. The results in Table 1a indicate that there was no difference in the outcome whether or not an exact copy of the test image was present in the database. The collective results indicate that CO1 can work as a robust image recognition system even when the database contains images of varying quality (images taken in good light and sharply in focus together with images produced in low light, softer focus). StripeSpotter automatically lets the user know the quality of the images used, every image in the database is classed as good, bad or ok, therefore providing the user with further information on the reliability of the results and whether or not to accept them. Every image, the test and all of the ranked images are visible to the viewer. The algorithm worked to accurately identify wings even if they were broken or clipped at the edges, as they were in many *T. brevipalpis*.

A larger database resulted in significantly more accurate identifications than using a smaller database - student’s t test result, with t>2.086, df = 20 at p=0.05; where p was < 0.05 in each case and a ‘difference in proportion test’, where the difference between the LDB and the SDB was statistically significant at the 95% level, except for the results in **bold**^*^ (no difference), Table 1. This tendency was also reflected in the values obtained when images further down the ranks were considered, up to rank 5 and 10; the larger database resulted in greater numbers of accurate identifications (Table 1). The trend continued to be reflected when the values for the mode and median were considered, both from early on in the ranking system (ranks 1&2) and further down the ranks (up to ranks 5 and 10); using the larger database resulted in higher modal and median values than using a smaller database (Table 2, larger numbers of accurate identifications further down the ranks).

The hind wings of *Bombus* sub species produced higher numbers of accurate identifications than the fore wings, indicating the importance of the hind wings when trying to separate sub species. It is speculated that CO1 may be taking the speckling on the hind wings into account, something which the human eye would find difficult to categorise and should be investigated further. CO1 did not confuse the two *Anopheles* species, *A. gambiae* and *A. stephensi* (Table 1, Figure 1). When these species are distinguished using traditional keys and the human eye, the dark and pale scale patterns on the upper part of the wings are generally used as distinguishing features and CO1 successfully separated these two very important vectors of malaria with a high degree of accuracy. Further testing of many more anopheline species needs to be carried out to determine if CO1 can successfully separate other anopheline species when present in the database. However, when testing species separated by small differences, CO1 should be carefully employed, good images (‘’good’’ as indicated by the software and by eye) of both sexes in sufficient quantities should be present in the database and the test image used should also be of good quality.

The sexes were differentiated well in the majority of species. In mosquitoes, this could be due to CO1 picking up differences in the numbers of scales present on a female wing (much greater) compared to males (most evident on the last vein, marked by a red square, *A. gambiae*, Figure 1). In bees, this could be due to differences in the size of the wings – male wings are much larger than females in *A. mellifera*. Male hind wings of *A. mellifera* were recognised accurately 100% of the time, not just at rank 1 & 2, but also successively up to rank 30 - entire rows from rank 1 to rank 30 were accurately identified male hind wings (30 male hind wings present in the database).

Comparing the results from the third database (Table 3) with the appropriate results ‘B’ from the large database test (Table 1a) indicated that there was no significant difference in results. The 3^rd^ database, Table 3, comprised of 30 good images (‘’good’’ as indicated by the software and by eye) of each of the species, hence using only good images in the database in sufficient numbers (30 here, compared to the 12 good images of each species in the smaller database) and also testing with good images resulted in similar outcomes as when the large database, with mixed quality images, was used. Therefore in any test, it is important to know the composition of what is in the database and the quality and quantity of the database images. This information is automatically formulated within StripeSpotter when the database is constructed and can be easily accessed.

The cost values indicated that if an exact copy of a test image was in the database, it would invariably be retrieved at rank 1 and the average value would be 0.999 (Table 3). If an image was tested that was far removed/completely different from wing images in the database, such as images of houses, faces, cars, trees, the average cost value was 0.92 (rounded up) hence any value at or below this was not a wing image. An average cost value of 0.9849 was obtained when wings were tested using a database which did not contain any wing images at all of the test species, here the database consisted of wing images from only one species (A. *stephensi*). Hence, any cost value at or below 0.9849 would be an indication that images of the test species were not present in the database being used. An average cost value of 0.99653 was obtained when images of the test species were present in the database (but not an identical copy of the test). This indicates that any cost value at rank 1, at or below 0.995 (to allow for a wide margin of error) would need to be further scrutinised. If cost values are to be used for any purpose, then they should be calculated for each database created, as database content can vary.

Regardless of any cost value, or rank 1, up to rank 5 or until rank 10 values, it should be noted that the test image and all of the retrieved ranked images (up to rank 100) can all be seen by eye when using the software and any image could be rejected as not being accurately identified at rank 1 if it appeared to be incorrect and warranted further scrutiny. It is recommended that any insect identification web site using software similar to CO1 also carried further images of other body characteristics and salient features of the insect anatomy of note, to aid greater certainty of accurate species identification. Where possible, the species present in the database should have salient features of the species specified on the web site, so that all rank 1 and up to rank 10 identifications can be checked for other features of the anatomy that matched the test species and the retrieved ranked results. For example if a rank 1 image was that of *A. aegypti*, the additional information on the web site should state that this species had scale markings on the thorax in the shape of a lyre and the test species should be checked for all such salient features. Image recognition software such as CO1, is best used in tandem with a description of easily recognised, salient, anatomical features of all the species in the database. This is because although the levels of accurate identifications are very high, they are not 100% in every case/species. Using additional easily recognizable morphological information about all of the species in the database as well as rank 1 and up to rank 5 and 10 results, plus the cost values of rank 1 images, would allow for greater assurance of the identification.

CO1 can be a powerful aid and useful tool in scientific studies where large numbers of different insect species need to be sorted and the identification character need not be restricted to the wings, any part of the insect anatomy can be utilised, provided the images are all consistently aligned - CO1 is adept at recognising curves, so consistent wing alignment is very important. Preparing samples for identification using CO1 takes the same time as preparing them for traditional identification. However there is no need to go through identification keys from a manual, or to have advanced taxonomical knowledge of every species. Software such as CO1, especially if it can be adapted for insect identification, is a useful tool for moving identification of insect species from a physical page in a book (niche information, not always easily or widely accessible) to electronic identification easily accessible to everyone.

Previous studies have shown that image recognition of insects has important applications in industry for the identification of pests of stored food and similarly in agriculture to identify quickly, pests of growing food crops. This study has shown that image recognition can also be an aid in the identification of disease bearing insects such as mosquitoes. Its use in the identification of beneficial insects such as bees (ABIS) has already been intensely studied (Al-Saqer et al 2011; An Lu 2010; Hall 2011; Jing Dai 2006; Lamprecht 2010; Wang et al 2012; Yang et al 2010; Zhao et al 2009; and Zhou et al 1985). However, very few insect identification web sites utilise image recognition software as part of the identification process. In contrast to the identification of trees, where species recognition using images of tree leaves is now being established and freely available on the leafsnap website (leafsnap.com, still being developed) and the leafsnap UK App (available now and free to use on iTunes).

The use of CO1 in citizen science studies (Davies et al 2012; Gura Trisha 2013), where lay members of the public contribute to scientific studies, is not straightforward in the case of insect identification where the wings are used. This is because unlike zebras (Lahiri et al 2011a & b), whale sharks (Davies et al 2012) or any other large animal where Citizen Science has led to a body of scientific contribution by the public; in the case of insect wings such as mosquito wings, the size of the organism and the fact of having to dissect out the wings, could present difficulties. However, this need not be a major limitation as technological advances mean that magnifying attachments are available for many modern cell/mobile phones and technical issues such as these can be overcome. Not every species of insect is as small as a mosquito, bee wings which are larger can be photographed well using a macro lens and the wing need not be dissected. Simply placing a white piece of paper in between the resting wing and the body and taking a photo with overhead lighting from a shaded lamp would suffice to capture the necessary vein detail. Due to their larger size, it is also possible to place the wings of bees between 2 microscope slides and take a good photograph without dissecting them from the body. Methods of obtaining good images without dissecting the wings need to be investigated further. One of the advantages of this study is that the insect is dead and therefore does not move, affording the opportunity to obtain decent images compared to a larger, moving object some distance away such as Zebras and Whales. Image capturing technology is advancing rapidly and it should be possible to capture even smaller insect wings with a simple and inexpensive USB digital microscope attached to a laptop and take excellent photographs. The capabilities of the dedicated citizen scientist should not be underestimated and in cases where obtaining images from larger insects is relatively easy, citizen science contributions should be considered, especially if CO1 can be tailored for insect wing identification and available as an Application (App).

Such an App would be an invaluable aid to researchers in the field, for example where workers needed to identify different insect species from traps. It would also be an aid to anyone involved in the production of food and wishing to identify insect pests that may be attacking their crop. The applications are numerous and producing stock images of insects which are then uploaded into a ‘World Wide Database’ of insect images should become a primary aim in our digital age, where all known insect images uploaded can be made available to everyone for image recognition, as insects impact greatly on human health and food production. This is a realistic goal as entomologists and others working with or concerned with insects can be invited to upload ordered images onto this ‘Global Database of Insect Images’.

## Conclusions

The ability of CO1 on the StripeSpotter platform to identify insect wings was assessed with promising results.

In conjunction with other salient features of all the species present in the database, as well as the results from the ranked images, CO1 can be reliably employed in the accurate identification of insect species as long as it is used correctly. It is suggested that image recognition software be utilised on insect identification web sites especially for species where wing identification is the norm.

Modern image recognition software such as CO1 can be ideal tools for use on insect identification websites by the general public (the citizen scientist). It can be just as useful in scientific research where speedy and accurate sorting of large numbers of insect species is required.

## Acknowledgements

This work would not have been possible without the generous donation of dead insects. Warm thanks are due to Peter Bowbrick (Surrey Beekeeper’s Association) for the supply of honeybees. To Dr Mark Rowland and Shahida Begum (London School of Hygiene and Tropical Medicine) for the mosquito species and to Gemma Baron and Prof Mark Brown (Royal Holloway College, Surrey) for the bee and bumblebee species – many thanks. Communication with the founders of StripeSpotter, Dr Mayank Lahiri (Google), Prof Tanya Berger-Wolf (University of Illinois, US) and of HotSpotter, Prof Charles Stewart (Rensselaer Polytechnic Institute, US) and Prof Daniel I Rubenstein (Princeton University) is recognized. Finally, encouragement from the Daphne Jackson Trust is acknowledged.

